# TLR8-Activating miR-146a-3p is an Intermediate Signal Contributing to Fetal Membrane Inflammation in Response to Bacterial LPS

**DOI:** 10.1101/2023.11.14.567042

**Authors:** Hanah M. Georges, Caterina Cassin, Mancy Tong, Vikki M. Abrahams

**Author notes:** **Corresponding author:** Vikki M Abrahams, PhD. Department of Obstetrics, Gynecology & Reproductive Sciences, Yale School of Medicine, 310 Cedar Street, LSOG 305C, New Haven, CT 06510, USA.

## Abstract

Preterm birth is the largest contributor to neonatal morbidity and is often associated with chorioamnionitis, defined as inflammation/infection of the fetal membranes (FMs). Chorioamnionitis is characterized by neutrophil infiltration of the FMs and is associated with elevated levels of the neutrophil chemoattractant, interleukin (IL)-8, and the proinflammatory cytokine, IL-1β. While FMs can respond to infections through innate immune sensors, such as Toll-like receptors (TLRs), the downstream mechanisms by which chorioamnionitis arises are not fully understood. A novel group of non-classical microRNAs (miR-21a, miR-29a, miR-146a-3p, Let-7b) function as endogenous danger signals by activating the ssRNA viral sensors TLR7 and TLR8. In this study, the pro-inflammatory roles of TLR7/TLR8-activating miRs were examined as mediators of FM inflammation in response to bacterial lipopolysaccharide (LPS) using an *in vitro* human FM explant system, an *in vivo* mouse model of pregnancy, and human clinical samples. Following LPS exposure, miR-146a-3p was significantly increased in both human FM explants and in wildtype mouse FMs. Expression of miR-146a-3p was also significantly elevated in FMs from women with preterm birth and chorioamnionitis. FM IL-8 and inflammasome-mediated IL-1β production in response to LPS was dependent on miR-146a-3p and TLR8, downstream of TLR4 activation. In wildtype mice, LPS exposure increased FM IL-8 and IL-1β production and induced preterm birth. In *TLR7^−/−^/TLR8^−/−^*mice, LPS exposure was able to initiate, but not sustain preterm birth, and FM inflammation was reduced. Together, we demonstrate a novel signaling mechanism at the maternal-fetal interface in which TLR8-activating miR-146a-3p acts as an intermediate danger signal to drive FM inflammasome-dependent and -independent mechanisms of inflammation and thus, may play a role in chorioamnionitis and subsequent preterm birth.

## Introduction

Preterm birth, defined as parturition prior to 37 weeks of gestation, afflicts ∼10% of U.S. pregnancies (1), and can have lifelong impacts on the health of the offspring, contributing to 75% of neonatal mortality and 50% of long-term neurological disorders in children (2). Chorioamnionitis, which is defined as fetal membrane (FM) inflammation characterized by neutrophil infiltration, is a significant cause of preterm birth and is often associated with bacterial infections. However, most women with chorioamnionitis fail to present signs of infection or inflammation, making diagnosis and treatment particularly challenging (3). Despite continued research, the mechanisms underlying chorioamnionitis and its contribution to preterm birth remain unclear, presenting a significant clinical problem for maternal-fetal medicine.

Inflammation is a critical component of pregnancy success; it is needed for proper implantation, parturition/labor, and to protect the fetus against invading pathogens. Upon the onset of normal term labor, an inflammatory process is activated which allows for the rupture of the FMs, cervical ripening, and a healthy delivery. However, untimely induction of inflammation, in response to invading pathogens, can prematurely activate the inflammatory processes of labor, contributing to preterm birth. Like many tissues, FMs express innate immune pathogen recognition receptors (PRRs), such as Toll-like receptors (TLRs), which sense and respond to infectious components (4, 5). When activated, these receptors contribute to an inflammatory response by inducing the production of pro-inflammatory chemokines and cytokines. In human FMs, activation of TLRs by bacterial components can lead to the production of the neutrophilic chemokine, IL-8, and the pro-inflammatory cytokine and mediator of tissue injury, IL-1β (4, 6); both established markers of preterm birth (7, 8). Despite FMs possessing the ability to mount an innate immune response, the balance between protecting the mother and fetus, and triggering preterm birth is delicate and there is still much about the molecular mechanisms that are unknown. While TLRs can directly activate inflammatory signaling pathways in response to an infection, there is potential for more complex modulation, regulation, and fine-tuning of these signaling processes and the type of responses generated (9). One regulatory mechanism is the requirement for two or more PRRs to be activated in combination or sequentially (9). This is well represented by the inflammasome where signal one is typically activation of a PRR inducing pro-IL-1β expression, followed by a second signal that activates the inflammasome-associated Nod-like receptor (NLR) or sensor for subsequent IL-1β processing and secretion (10). This second signal may be either an infectious trigger or a host-derived danger signal or damage-associated molecular pattern (DAMP) (10). For example, the DAMP, uric acid, is an activator of the NLPR3 inflammasome (11) and plays a role in mediating FM IL-1β production in response to lipopolysaccharide (LPS) (6). It is likely that there are additional DAMPs that can modulate inflammasome-dependent and inflammasome-independent mechanisms of inflammation at the maternal-fetal interface that have yet to be explored.

Classically, microRNAs (miRs) are small noncoding RNAs that regulate protein expression through post-transcriptional interactions with mRNA (12). However, a novel group of miRs, with specific GU rich regions, act non-classically as DAMPs by activating endosomal TLR7 and/or TLR8 that typically detect viral ssRNA (13). miR-21a, miR-29a, miR-146a-3p and Let-7b are ligands for TLR7 and/or TLR8 that can trigger an inflammatory response (14–16). Since TLR7 and TLR8 are expressed by human FMs (4), we questioned the pro-inflammatory roles of these miRs and their potential to mediate the production of critical inflammatory factors known to contribute to chorioamnionitis and preterm birth. We hypothesized that in response to a bacterial stimuli, TLR7/TLR8-activating miRs act sequentially downstream of an initial sensor to mediate FM inflammation, specifically chemotactic IL-8 and inflammasome mediated IL-1β.

Using an established *in vitro* human FM explant model, an *in vivo* mouse model of pregnancy, and *in vivo* human clinical samples, we demonstrate, for the first time, a pro-inflammatory role for FM miR-146a-3p through TLR8 activation, which is dependent upon a prior primary signal from TLR4 activation by bacterial LPS. Furthermore, TLR8-activating miR-146a-3p is elevated in FMs from women with preterm birth and chorioamnionitis, and using a mouse model we report a role for TLR8 in contributing to FM inflammation and preterm birth.

## Materials and Methods

### Patient Samples

Human FM tissue collection was approved by the Yale University’s Human Research Protection Program (IRB# 0607001625) and was facilitated by the Yale University Reproductive Sciences (YURS) Biobank (IRB# 1309012696) with patient consent. For all *in vitro* explant experiments, human FMs were collected from uncomplicated term pregnancies (38-41 weeks gestation) delivered by scheduled caesarean section, without evidence of labor or infection, and were transferred to the laboratory immediately after delivery. For the analysis of FMs from women with different pregnancy outcomes, tissues were collected at the time of delivery and were immediately washed, snap frozen, and stored at −80°C prior to analysis. These banked human FMs were accessed from the following groups: 1. Uncomplicated pregnancies delivered at term (38-41 weeks) vaginally and with labor (Term + Labor) [n=9]; 2. Uncomplicated pregnancies delivered at term (38-41 weeks) by cesarean-section without labor (Term - Labor) [n=13]; 3. Preterm birth with chorioamnionitis (PTB + CAM) [n=7]. These were women spontaneously delivered prematurely (<37 weeks) with preterm premature rupture of membranes (PPROM) who were positive for chorioamnionitis as defined by infection and/or inflammation determined by an amniotic culture and/or histological examination; 4. Preterm birth without chorioamnionitis (PTB - CAM) [n=7]: These were women spontaneously delivered prematurely (<37 weeks) for obstetrical indications not associated with preeclampsia or chorioamnionitis. These are classed negative for infection and/or inflammation, determined by an amniotic culture and/or histological exam; 5. Preterm birth with preeclampsia (PTB +PE) [n=11]. These were women who were delivered prematurely (<37 weeks) due to preeclampsia. Patient demographics and clinical information are summarized in Table 1.

**Table 1.** Clinical and demographic characteristics of fetal membrane donors for miR-146a-3p expression analysis. Data were analyzed using chi-squared tests.

|  | **Term + Labor** | **Term - Labor** | **PTB + CAM** | **PTB - CAM** | **PTB + PE** | **p value** |
| --- | --- | --- | --- | --- | --- | --- |
| **N** | 9 | 13 | 7 | 7 | 11 |  |
| **Maternal Age, Years, Median, (IQR)** | 32 (31-34) | 33 (32-35) | 33 (30.5-37.5) | 32 (27.5-35) | 34 (30.5-38.5) | 0.369 |
| **Race, n (%)** |  |  |  |  |  |  |
| **Caucasian** | 7 (77.8) | 13 (100) | 4 (57.1) | 2 (28.6) | 5 (45.5) |  |
| **Black / African American** | 0 (0) | 0 (0) | 0 (0) | 2 (29) | 3 (27) |  |
| **Asian** | 1 (11) | 0 (0) | 1 (14.3) | 1 (14.3) | 0 (0) |  |
| **Hispanic** | 1 (11) | 0 (0) | 2 (29) | 2 (29) | 1 (9) |  |
| **Maternal BMI, Median, (IQR)** | 21.8 (20.3-24.2) | 30.6 (27.1-33.7) | 31.8 (29.1-34.5) | 30.1 (26.7-37.1) | 24.8 (23.8-33.8) | 0.007 |
| **Primiparity? N, (%)** | 6 (67) | 3 (23) | 4 (57) | 6 (86) | 4 (36) | 0.054 |
| **Gestational Age at Delivery, median weeks, (IQR)** | 40.3 (40.1-40.7) | 39.42 (39.1-39.4) | 30.4 (26.8-34.7) | 34 (34.0-34.7) | 34 (34.0-36.2) | 0.0001 |
| **Birthweight, Median Grams, (IQR)** | 3630 (3580-3950) | 3870 (3450-4310) | 1430 (975-2300) | 2130 (1725-2280) | 2190 (1880-2740) | 0.0001 |
| **Male Infants, N (%)** | 6 (66.7) | 9 (69.2) | 2 (28.6) | 3 (42.9) | 6 (54.5) | 0.412 |

### Human fetal membrane explants

Human FM explants were prepared as previously described (4, 5, 17, 18). In brief, with both the chorion and amnion layers intact, FMs were biopsied using a 6mm punch. Biopsies were placed into 0.4μm cell culture inserts (BD Falcon, Franklin Lakes, NJ) in a 24 well plate with 1mL of Dulbecco modified eagle medium (DMEM; Gibco, Grand Island, NY) supplemented with 10% fetal bovine serum (Hyclone, Logan, UT) and 1% penicillin streptomycin (Gibco, Grand Island, NY). The next day, media was replaced with serum-free OptiMeM (Gibco, Grand Island, NY), rested for 3 hours, then treated as described below.

### Fetal membrane explant treatments and transfections

Human FM explants were treated with either a no treatment (NT) media control or 100ng/ml LPS isolated from *Escherichia coli* O111:B4 (Sigma-Aldrich, St Louis, MI) for 48 hours. For inhibitor studies, FMs were pretreated for 1 hour with or without either the TLR4 antagonist, LPS-RS (10μg/ml; Invivogen, San Diego, CA), the TLR7 inhibitor, IRS661 (5′- TGCTTGCAAGCTTGCAAGCA-3’; made endotoxin-free by the Keck Core, Yale University) (19–21), or the TLR8 inhibitor, CU-CPT9a (10μM; Invivogen, San Diego, CA) before subsequently treating with or without LPS. To test the efficacy of the TLR7 inhibitor, IRS661, HEK-TLR7/NFκB reporter cells (Invivogen, San Diego, CA) were treated overnight with or without or the TLR7 agonist, Imiquimod (R837; Invivogen, San Diego, CA) at 5μg/ml in the presence of media or IRS661 (5μM). Optical densities were read at 655nm. To assess miR-146a-3p function, FM explants were transfected using siPORT NeoFX (Invitrogen, Waltham, MA) with either a MirVana miR146a-3p mimic (PM13059; ThermoFisher Scientific, Waltham, MA) at 0.01nM for 24 hours, a MirVana anti-miR146a-3p inhibitor (MH13059; ThermoFisher Scientific, Waltham, MA) at 100nM for 24 hours, or the appropriate miR mimic or inhibitor scramble control at 100nM (ThermoFisher Scientific, Waltham, MA). Transfections were confirmed by reverse transcription quantitative polymerase chain reaction (RT-qPCR) for miR-146a-3p. From all experiments, supernatants were collected on ice and FMs were snap frozen on dry ice. Both sample types were stored at −80°C prior to subsequent analysis.

### Mouse studies

All animal studies were conducted in accordance with NIH guide for the care and use of laboratory animals and approved by Yale University’s institutional animal care and use committee (#2022-11589). This study utilized two mouse strains at 8-12 weeks of age: *C57BL/6* wildtype mice from Jackson Laboratories (Bar Harbor, ME) and *TLR7^−/−^/TLR8^−/−^* mice on a *C57BL/6* background which were a generous gift from Professor Richard Flavell (22). Due to the presence of autoimmune disease in single *TLR8^−/−^* single knockout mice (23) which could impact our studies, we used *TLR7^−/−^/TLR8^−/−^* double knockout mice which do not exhibit autoimmune-like symptoms (23). In both wildtype and *TLR7^−/−^ /TLR8^−/−^* mice, timed pregnant females were injected intraperitoneally (ip) with 100μl of PBS or LPS (50μg) on day E15.5 as previously described (18). Mice were either monitored for preterm birth for 48 hours post injection or sacrificed 6 hours post injection when FM tissue samples were dissected from the placental unit as previously described (17, 24), pooled from each dam, snap frozen, and stored at −80°C prior to subsequent analysis.

### RNA isolation and RT-qPCR

Human and mouse FMs were homogenized using a beadbug microtube homogenizer with microtubes pre-filled with high impact zirconium beads (Benchmark Scientific; Sayreville, NJ) as previously described (6). RNA was extracted via TRIzol (ThermoFisher Scientific; Waltham, MA) according to manufacturer’s protocol (17, 24). RNA concentrations were measured using a NanoDrop 2000 Microvolume Spectrophotometer (ThermoFisher Scientific; Waltham, MA). Human and mouse FM tissue expression of miR-21a, miR-29a, miR-146a-3p, and Let-7b were measured using the Taqman MicroRNA Assay (Life Technologies; Carlsbad, CA) according to manufacturer’s protocols for reverse transcription and qPCR. For miR-146a-3p, a pre-amplification step was included prior to performing PCR using the Taqman PreAmp master mix kit (Life Technologies; Calrsbad, CA). Small nuclear RNA U6 was used as a reference gene for the normalization of miR expression. Mouse FM expression of *IL1B* and *KC* was measured by RT-qPCR using the SuperScript II kit (Invitrogen; Waltham, MA) and the Kapa SYBR Fast qPCR kit (Sigma-Aldrich; St. Louis, MO) and normalized to *GAPDH*. Primer sequences have been previously reported (17, 25). RT-qPCR data was analyzed using the 2^−ΔΔCT^ method and reported as either fold change or relative abundance (RA).

### FM supernatant analysis

Supernatants collected from human FM explant experiments were measured for the chemokine IL-8 and cytokine IL-1β by ELISA (R&D Systems, Minneapolis, MN). FM uric acid secretion was measured using a colorimetric assay (Sigma-Aldrich; St. Louis, MO).

### Statistical analysis

Experiments were performed three or more times. The number of independent experiments that data were pooled from are indicated in the figure legends as “n=”. Data are reported as mean ± SEM with statistical significance defined as *p*<0.05. Statistical testing were performed using Prism Software (Graphpad, La Jolla, CA). For normally distributed data, significance was determined using either one-way analysis of variance (ANOVA) for multiple comparisons or a t-test. For data not normally distributed, significance was determined using a non-parametric multiple comparison test or the Wilcoxon matched-pairs signed rank test. Additional statistical analyses are specified in Table legends.

## Results

### LPS induces inflammation and increases miR-146a-3p expression in human FM explants in a TLR4 dependent manner

Following LPS treatment, both chemotactic IL-8 and pro-inflammatory IL-1β secretion were significantly increased in human FM explant cultures (Figure 1A), recapitulating previous reports (4). Human FM expression levels of the TLR7/TLR8-activating miRs, miR-21a, miR-29a, and Let-7b were unchanged in response to LPS stimulation (Figure 1B). However, expression of TLR8-activating miR-146a-3p (16) was significantly increased in LPS treated FMs compared to the no treatment (NT) control by 2.1 fold ± 0.4 (Figure 1B). The presence of the TLR4 inhibitor, LPS-RS, significantly reduced FM secretion of IL-8 and IL-1β in response to LPS by 35.2 ± 9.1% and 58.3 ± 15.0%, respectively, when compared to the uninhibited (media) LPS control (Figure 1C). LPS-induced FM miR-146a-3p expression was also significantly decreased by 30.8 ± 17.3% in the presence of LPS-RS compared to the uninhibited (media) LPS control (Figure 1C).

**Figure 1.**
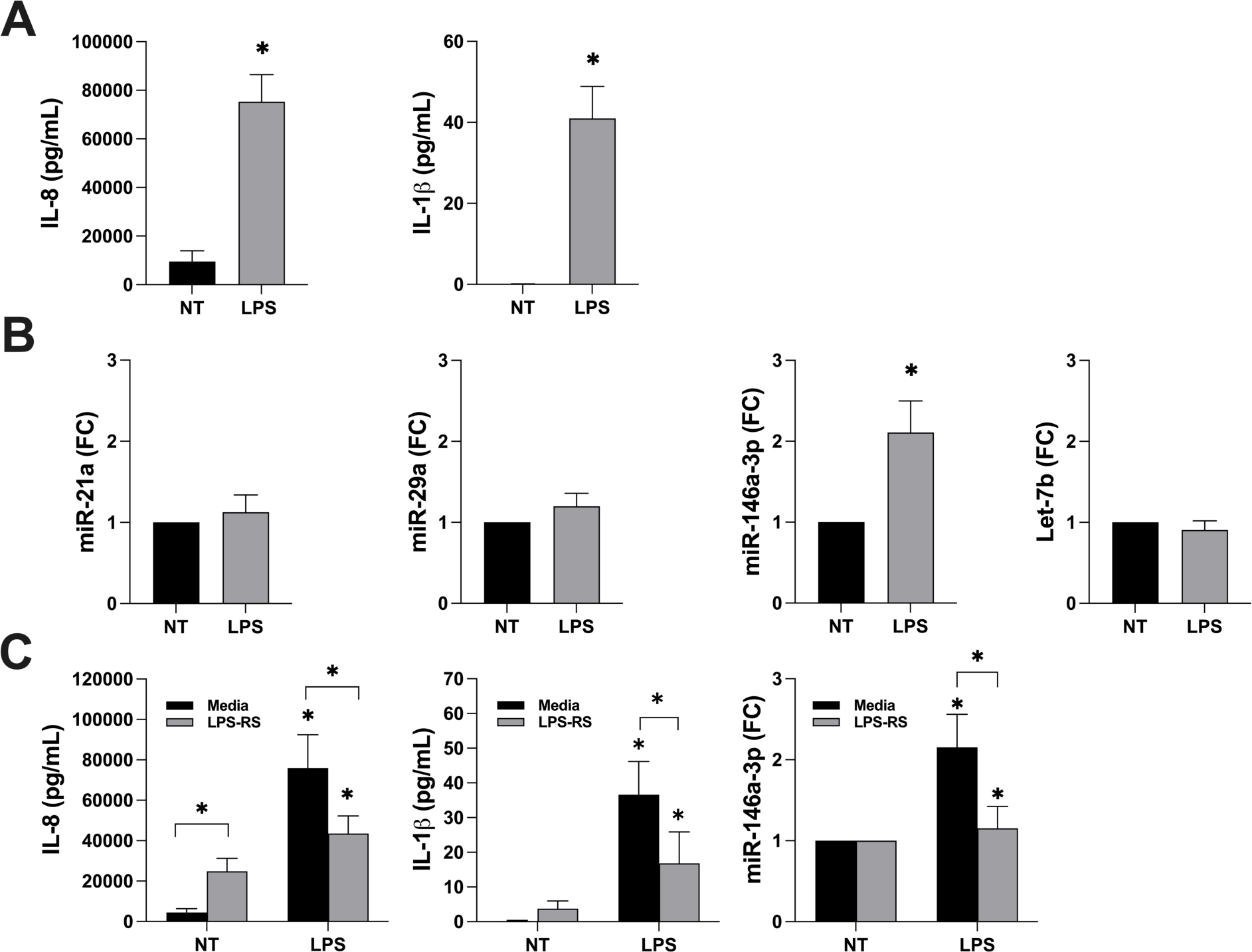
LPS induces inflammation and increases miR-146a-3p expression in human FM explants in a TLR4 dependent manner. (A-B) Human FM explants were treated with no treatment (NT) or LPS for 48 hrs after which: A) IL-8 and IL-1β secretion were measured by ELISA (n=13); and B) miR-21a, miR-29a, miR-146a-3p, and Let-7b were measured by RT-qPCR and expressed as fold change (FC) (n=13). \**p*<0.05 *vs.* the NT. (C) Human FM explants were treated with no treatment (NT) or LPS in the presence of media or the TLR4 inhibitor, LPS-RS. After 48 hrs IL-8 and IL-1β secretion were measured by ELISA (n=9), and miR-146a-3p was measured by RT-qPCR and expressed as fold change (FC) (n=9). \**p*<0.05 *vs.* the NT control.

### LPS-induced secretion of IL-8 and IL-1β by human FMs requires TLR8 activation

Since expression of miR-146a-3p was increased in FMs exposed to LPS, TLR7 and TLR8 were investigated as subsequent secondary signals for LPS/TLR4-induced FM inflammation. Human FM explants were pretreated with media, the TLR7 inhibitor, IRS661, or the TLR8 inhibitor, CU-CPT9a, followed by LPS treatment. Under all conditions, LPS significantly increased FM secretion of IL-8 and IL-1β when compared to the NT controls (Figure 2A). In the presence of the TLR8 inhibitor, CU-CPT9a, LPS-induced IL-8 and IL-1β secretion was significantly reduced by 42.7 ± 4.2% and 45.6 ± 10.6%, respectively, when compared to the uninhibited (media) LPS control (Figure 2A). In contrast, inhibition of TLR7 signaling by IRS661 had no effect on the ability of LPS to increase FM secretion of IL-8 and IL-1β (Figure 2A). Efficacy of IRS661 was tested in TLR7 reporter cells stimulated with the TLR7 agonist Imiquimod, and this confirmed its inhibitory function (Figure 2B). Since we previously reported that LPS induces human FM IL-1β secretion through endogenous uric acid activation of the NLRP3 inflammasome (6), we tested the regulation of FM uric acid by TLR8. As shown in Figure 2C, the presence of the TLR8 inhibitor, CU-CPT9a, significantly reduced FM uric acid levels under LPS conditions.

**Figure 2.**
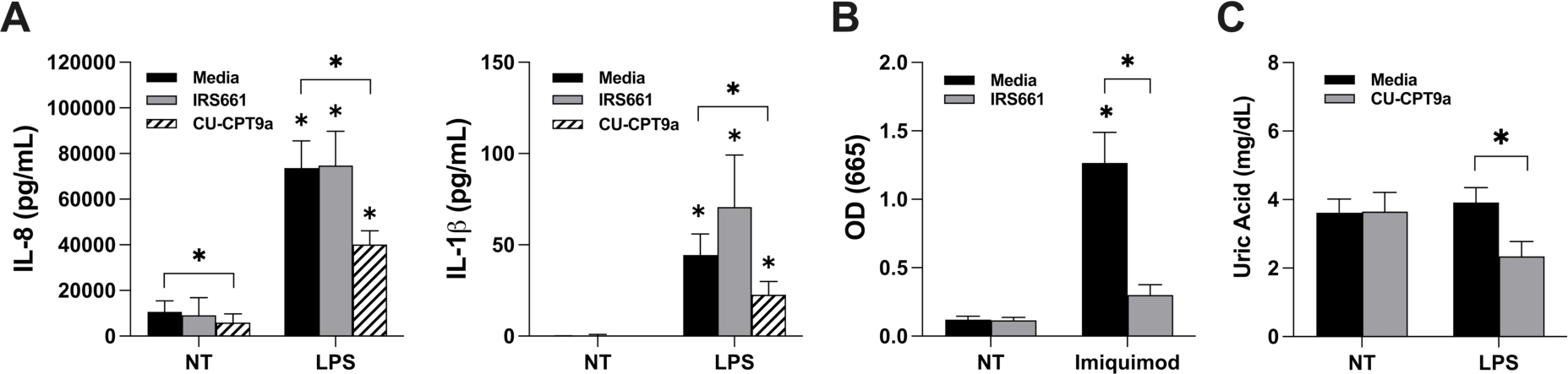
LPS-induced secretion of IL-8 and IL-1β by human FMs requires TLR8 activation. A) Human FM explants were treated with no treatment (NT) or LPS in the presence of media; the TLR7 inhibitor, IRS661; or the TLR8 inhibitor, CU-CPT9a (n=7-11). After 48 hrs, IL-8, IL-1β were measured. B) HEK-TLR7/NFκB reporter cells were treated with NT or the TLR7 agonist, Imiquimod (R837) in the presence of media or the TLR7 inhibitor, IRS661 after which optical densities (OD) were read at 655nm (n=3). C) Human FM explants were treated with no treatment (NT) or LPS in the presence of mediator the TLR8 inhibitor, CU-CPT9a (n=9). After 48 hrs, uric acid levels were measured. For all charts, \**p*<0.05 *vs.* the NT control unless otherwise indicated.

### miR-146a-3p mediates human FM inflammation in response to LPS

Given that LPS-induced FM inflammation is dependent upon the LPS sensor TLR4 and the ssRNA sensor TLR8, we sought to determine if LPS-induced miR-146a-3p was mediating this sequential signaling pathway. As shown in Figure 3, transfection of human FM explants with a specific anti-miR-146a-3p inhibitor significantly inhibited the ability of LPS to induce FM secretion of IL-1β by 32.5 ± 12.3% when compared to the scramble control. The anti-miR-146a-3p inhibitor did not affect the ability of LPS to induce IL-8 secretion (Figure 3). However, in the absence of LPS, transfection of human FM explants with a specific miR-146a-3p mimic elevated IL-8 secretion by 1.5 ± 0.3 fold and this was reduced to below baseline by the presence of the TLR8 inhibitor, although significance was not reached (Supplementary Figure 1). Transfection of human FMs with the miR-146a-3p mimic, however, did not elevate IL-1β secretion (Supplementary Figure 1).

**Figure 3.**
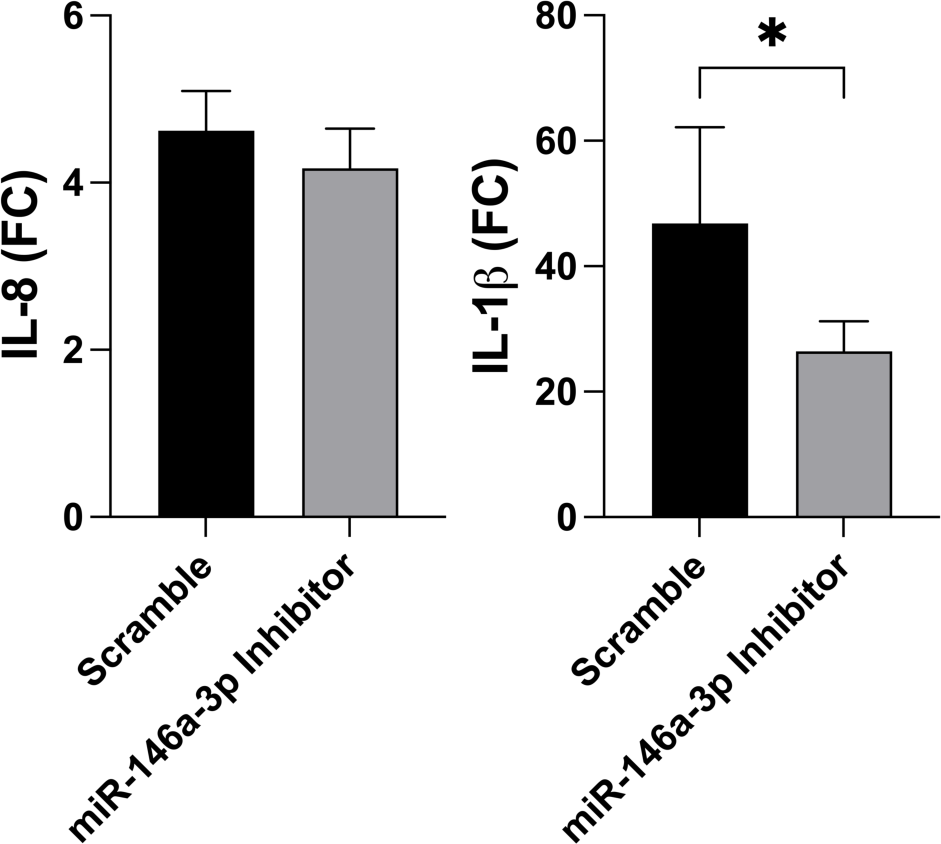
miR-146a-3p mediates human FM inflammation in response to LPS. Human FMs were transfected with either an anti-miR-146a-3p inhibitor or a scramble control and then treated with or without LPS. After 48 hrs, IL-8 and IL-1β secretion was measured. Data are shown as LPS induced fold change (FC) (n=18). \**p*<0.05 *vs.* the scramble control.

### FM miR-146a-3p is elevated in a mouse model of LPS-induced preterm birth

We next sought to validate our *in vitro* data using an established *in vivo* mouse model of LPS-induced preterm birth (26). Following ip administration of high dose LPS to wildtype mice, the rate of preterm birth was significantly higher at 80.0 ± 20.0% when compared to PBS treated control mice that displayed no preterm birth (*p*=0.048; Table 2). Following model validation, FMs were collected from PBS and LPS exposed mice 6 hours post injection. Similar to our findings in human FMs *in vitro*, mouse FM *miR-146a-3p* expression was significantly elevated in response to LPS when compared to the PBS controls, while FM expression of *miR-21a*, *miR-29a* and *Let-7b* were unaffected (Figure 4A).

**Figure 4.**
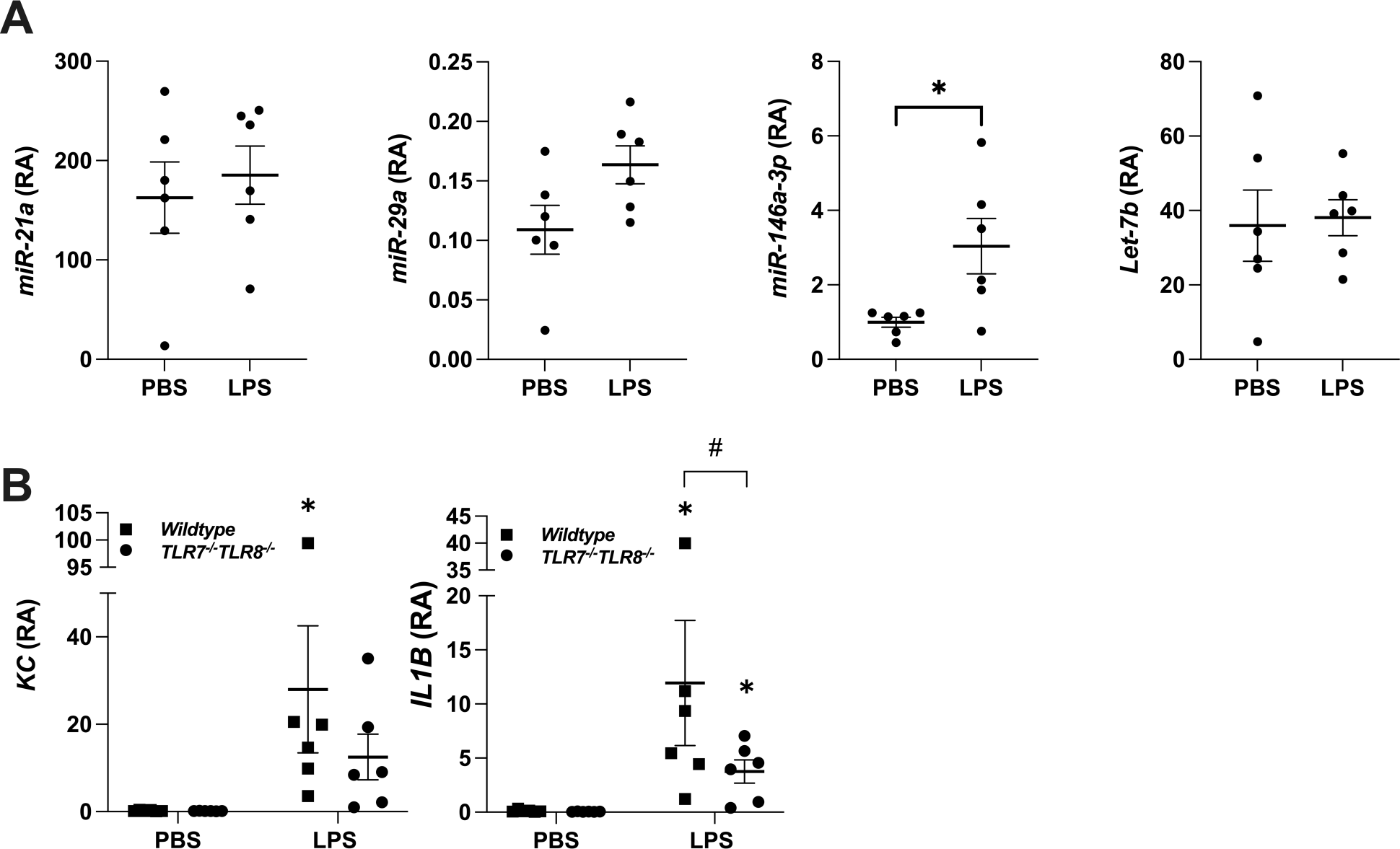
FM miR-146a-3p is elevated in a mouse model of LPS-induced preterm birth. (A) Wildtype pregnant mice were administered PBS (n=6) or LPS (n=6) on E15.5. 6 hrs later dams were sacrificed, FM were collected and analyzed by RT-qPCR for *miR-21a, miR-29a, miR-146a-3p, and Let-7b* expression. Data are expressed as relative abundance (RA); \**p*<0.05 *vs.* the PBS controls. (B) Wildtype and *TLR7^−/−^/TLR8^−/−^* pregnant mice were administered PBS (n=6) or LPS (n=6) on E15.5. 6 hrs later dams were sacrificed, FM were collected and analyzed by RT-qPCR for *KC* and *IL1B* expression. Data are expressed as relative abundance (RA); \**p*<0.05 *vs.* the PBS controls; and #*p*<0.09 as indicated.

**Table 2.** Preterm birth (PTB) percentage in wildtype (n=5) and *TLR7^−/−^/TLR8^−/−^* (n=6) mice exposed to either PBS or LPS. Comparisons were made within genotypes, PBS *vs.* LPS treatments. Data were analyzed using the fisher’s exact test.

| *Genotype of Dams* | *Treatment* | *N of Dams* | *Full PTB* | *Partial PTB* | *No PTB* | *p value* |
| --- | --- | --- | --- | --- | --- | --- |
| **Wildtype** | PBS | 5 | 0 (0%) | 0 (0%) | 5 (100%) | - |
| **Wildtype** | LPS | 5 | 4 (80%) | 0 (0%) | 1 (20%) | 0.048 |
| ***TLR7<sup>-/-</sup>/TLR8<sup>-/-</sup>*** | PBS | 6 | 0 (0%) | 0 (0%) | 6 (100%) | - |
| ***TLR7<sup>-/-</sup>/TLR8<sup>-/-</sup>*** | LPS | 6 | 0 (0%) | 6 (100%) | 0 (0%) | 0.002 |

### FM inflammation and preterm birth is initiated but not fully executed in pregnant TLR7^−/−^***/TLR8^−/−^ mice exposed to LPS***

Previous studies demonstrated that *TLR8^−/−^* single knockout mice develop autoimmune pathology (23), which are not present in *TLR7^−/−^/TLR8^−/−^* mice (22). With this in mind, as well as the potential for mouse TLR7 to function as a homologue to human TLR8 (13), *TLR7^−/−^/TLR8^−/−^* mice were used to determine the role of TLR7/TLR8 in FM inflammation and preterm birth in the presence of LPS. Interestingly, 100% of *TLR7^−/−^/TLR8^−/−^* mice treated with LPS exhibited partial preterm birth compared to 0% of *TLR7^−/−^/TLR8^−/−^* treated with PBS (*p*=0.002; Table 2). 12-15 hours following LPS administration, *TLR7^−/−^/TLR8^−/−^*mice presented with 1-2 fetuses expelled while the residual fetuses remained in the uterus. When FM inflammation was examined, *KC* (the mouse homologue of human IL-8) and *IL1B* expression were significantly increased in LPS exposed wildtype mice when compared to PBS treated control mice (Figure 4B). In contrast, FM *KC* was not significantly elevated in LPS exposed *TLR7^−/−^/TLR8^−/−^* mice when compared to the PBS controls (Figure 4B). While FM *IL1B* was significantly elevated in LPS exposed *TLR7^−/−^/TLR8^−/−^* mice when compared to the PBS controls, this trended towards lower expression compared to the wildtype mice (*p*=0.09, Figure 4B).

### Elevated human FM miR-146a-3p expression is associated with chorioamnionitis

To further examine the clinical relevance of our findings, human FMs were analyzed for miR-146a-3p expression from the following patient groups: Normal term deliveries with labor (Term + labor); normal term deliveries without labor (Term - labor); preterm birth deliveries with chorioamnionitis (PTB + CAM); preterm birth deliveries without chorioamnionitis (PTB - CAM); and preterm birth deliveries with preeclampsia (PTB + PE). FMs from women with preterm birth and chorioamnionitis expressed significantly higher levels of miR-146a-3p when compared to FMs from women with preterm birth without chorioamnionitis, with normal term controls with or without labor, and with preterm birth and preeclampsia (Figure 5).

**Figure 5.**
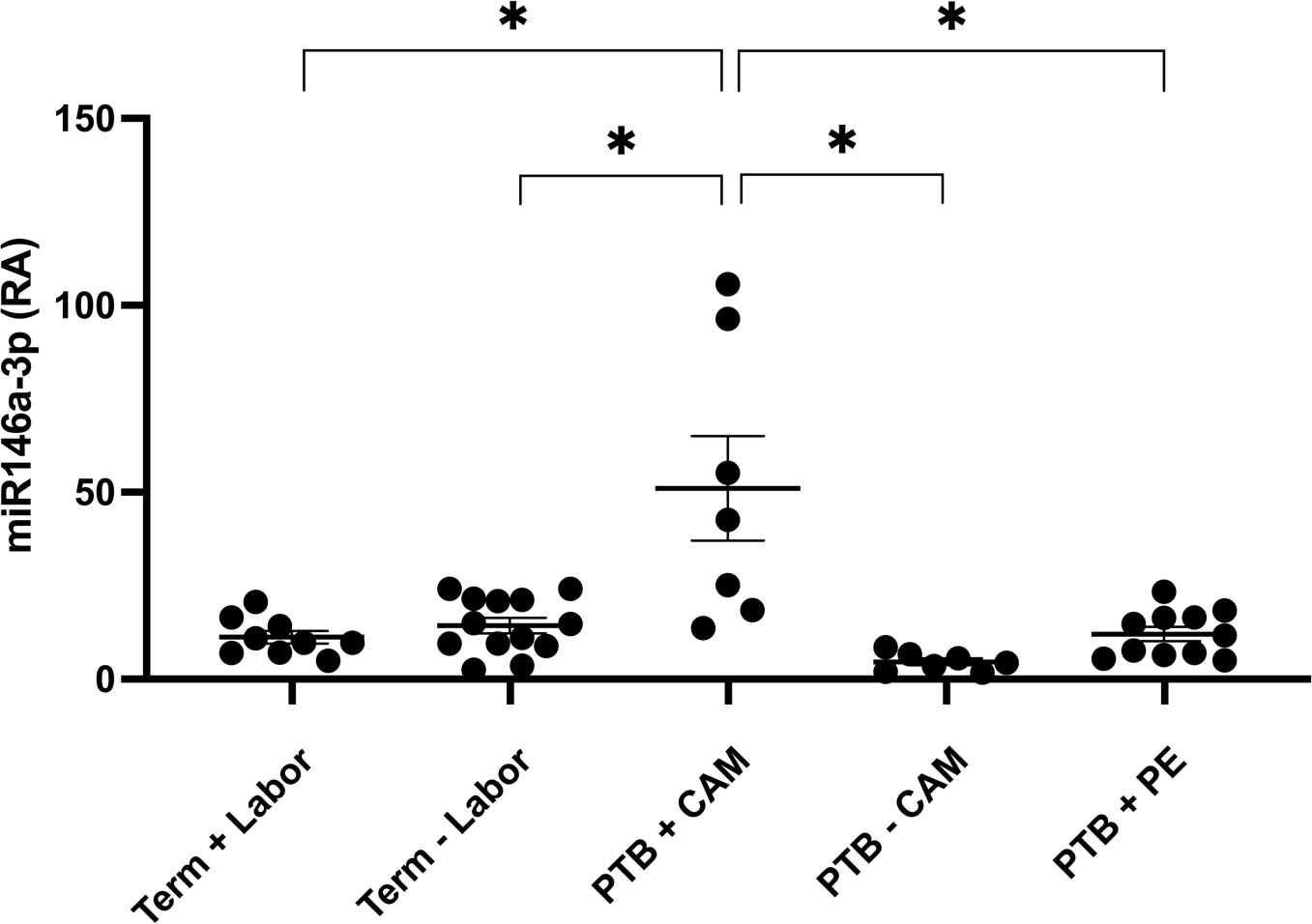
Elevated human FM miR-146a-3p expression is associated with chorioamnionitis. Expression of miR-146a-3p in FMs from the following clinical groups: Term with labor (Term + Labor, n=9); term without labor (Term - Labor, n=13); preterm birth with chorioamnionitis (PTB + CAM, n=7); preterm birth without chorioamnionitis (PTB - CAM, n=7); and preterm birth with preeclampsia (PTB + PE, n=11). Data are presented as relative abundance (RA), \**p*<0.05.

## Discussion

Despite advances in maternal and fetal care, preterm birth remains a worldwide health concern, especially as it remains the primary cause of neonatal death (27). Chorioamnionitis is seen in up to 70% of preterm birth cases and presents a significant threat to the health of the child (2). Additionally, the lifelong impacts of preterm birth on affected children are often overlooked, contributing to 50% of long-term neurological disorders and thus, a societal impact of $26.6 billion annually in the US (2, 27). Despite these statistics, a major factor in the persistence of chorioamnionitis and preterm birth as main contributors to neonatal morbidity is the difficulty in its diagnoses and lack of knowledge of the mechanisms of pathogenesis. The neutrophil recruiting chemokine, IL-8, and pro-inflammatory cytokine, IL-1β, are known markers of chorioamnionitis and preterm birth (7, 8, 28, 29), contributing to local inflammation in the placenta and FMs in response to a pathogenic insult. However, the innate immune signaling pathways involved in the secretion of these factors and their contributions to preterm birth are not fully understood. The current study sought to understand the fine regulation of FM inflammation in response to bacterial LPS through the examination of TLR7/TLR8-activating miRs, and their potential use as a diagnostic marker for chorioamnionitis. Herein, we report that bacterial LPS increases FM expression of miR-146a-3p which in turn, drives pro-inflammatory IL-8 and IL-1β production through activation of TLR8, contributing to preterm birth in a mouse model. Elevated FM miR-146a-3p is also associated with chorioamnionitis in humans.

This current study explored the non-canonical activation of TLR7 and TLR8 by miRs and their role in FM inflammation and preterm birth. Classically, miRs have been associated with translational regulation and mRNA degradation. However, recent studies revealed a unique family of miRs which act non-classically to agonize the endosomal ssRNA immune receptors, TLR7 and TLR8 (14, 15). miR-21a and miR-29a were found to create a pro-inflammatory and pro-metastatic state in a tumor microenvironment by activating TLR8 in human cells, activating TLR7 in mice, and inducing the expression of pro-inflammatory factors NFκB and TNFα (14). In another study, Let-7b was found to bind to neuronal TLR7 and activated the MyD88 signaling pathway leading to neurodegeneration (15). Let-7b was also able to increase NFκB expression following TLR7 activation in microglia (15). In a previous study by our group, miR-146a-3p was found to be upregulated in human placental trophoblast cells in a TLR4 dependent manner after exposure to antiphospholipid antibodies (aPL) (16). Moreover, aPL induced trophoblast miR-146a-3p promoted the secretion of IL-8 through activation of TLR8, suggesting TLR4 dependent miR-146a-3p further enhanced trophoblast inflammatory signaling through TLR8 in a sequential manner (16). With this discovery, we sought to understand miR-146a-3p and other TLR7/TLR8-activating miRs in a model of chorioamnionitis.

In the current study, we found miR-146a-3p expression was increased in human FM explants following TLR4 stimulation by bacterial LPS. Furthermore, secretion of pro-inflammatory IL-1β and the neutrophilic chemokine IL-8 was dependent on TLR8 activation downstream of TLR4 stimulation by LPS, corroborating previous work indicating non-canonical involvement of miR-146a-3p activation of TLR8 in placental inflammation (16). Of note, inhibition of TLR8 signaling only partially reduced the FM inflammatory response. Despite previous evidence of the pro-inflammatory actions of miR-21a, miR-29a, and Let-7b (14, 15), these miRs were not increased in human FMs in response to LPS in the current study, which may explain the lack of involvement of TLR7.

IL-1β is a highly regulated pro-inflammatory cytokine, typically requiring multiple signals for the secretion of its mature form. The two-hit hypothesis supports the concept that a first signal, usually from a pathogen, acts as a primer to either induce pro-IL-1β expression or increase expression of the inflammasome components, while a second signal, such as an additional pathogen or an endogenously produced DAMP, is required for inflammasome assembly and activation resulting in the cleavage of pro-IL-1β into its mature secreted form (10). We previously reported that bacterial LPS provides signal one and induces human FM IL-1β secretion through activation of the NLRP3 inflammasome by the production of an endogenous DAMP and NLRP3 agonist, uric acid, which acts as a secondary signal (6). In this current study we demonstrate the dependence of human FM uric acid and subsequent inflammasome-mediated IL-1β secretion in response to TLR4 activation by LPS on downstream TLR8 activation. Furthermore, inhibition of miR-146a-3p confirmed the role of this TLR8-activating miR in mediating human FM IL-1β secretion following LPS exposure. However, in the absence of the priming LPS signal, overexpression of miR146a-3p alone was insufficient to drive the FM IL-1β response. While we also demonstrated the dependence of human FM IL-8 secretion by LPS on TLR8 activation, inhibition of miR-146a-3p could not override this response, indicating additional downstream drivers of this chemokine. However, overexpression of miR-146a-3p did elevate FM IL-8 via TLR8 activation, although significance was not reached, again indicating additional signaling pathways or another TLR8-activating DAMP may be involved. Thus, miR-146a-3p-activation of TLR8 appears to act as an endogenous DAMP serving as a sequential signal for human FM secretion of IL-1β, and may contribute to FM IL-8 secretion, following LPS exposure.

We next went on to validate our human FM *in vitro* findings, *in vivo*. In animal models it has been shown that IL-1β, but not IL-8, is a known initiator of preterm labor (30). In mice, NLRP3-inflammasome-associated IL-1β contributes to preterm birth; NLRP3 inhibition reduced inflammation, delayed the onset of preterm birth, and reduced preterm birth rates (31). Nevertheless, IL-8 is still a pathological contributor as it is important for neutrophil chemotaxis and recruitment, a marker of chorioamnionitis, and *in vivo* models of chorioamnionitis have demonstrated elevated IL-8 in addition to IL-1β (28, 29). In our current study, in wildtype mice, LPS exposure resulted in preterm birth associated with increased FM expression of *miR-146a-3p*, *KC*, and *IL-1B*, which validated our *in vitro* human findings. In a non-pregnant system, a recent *in vivo* study also correlated miR-146a-3p with inflammation (32). However, this study was not associated with TLR7/TLR8 or inflammasome activation. Interestingly, in our experiments, the deletion of TLR7 and TLR8 in mice did not completely inhibit the initiation of preterm birth following LPS stimulation, but rather, arrested fetal expulsion, and this was associated with a significantly reduced FM inflammatory response. This indicates, similarly to our *in vitro* findings, that downstream of TLR4 activation by LPS, miR-146a-3p activation of TLR8 partially drives the FM inflammatory response which is required to sustain, but not needed for initiation of preterm birth. Thus other inflammatory pathways may be involved in the initiation of preterm birth. Whether mouse TLR8 is involved or mouse TLR7 is acting as the homolog for human TLR8 (13) is unclear since we chose to avoid our data being confounded by the autoimmune phenotype in the *TLR8^−/−^* single knockout mice (23).

For our human *in vivo* clinical sample validation study, FMs were collected from several different groups of pregnancies; term with labor, term without labor, preterm birth with chorioamnionitis, preterm without chorioamnionitis, and preterm birth due to preeclampsia. Strikingly, miR-146a-3p was only increased in the preterm with chorioamnionitis group. This implicates the importance of FM miR-146a-3p to specific inflammation based adverse pregnancy outcome and its potential as a diagnostic marker for FM inflammation.

While inflammation at the maternal-fetal interface is known to contribute to preterm birth, the precise mechanisms involved are still poorly understood. We have demonstrated that downstream of TLR4 activation by bacterial LPS, miR-146a-3p acts a novel danger signal intermediate for sequential activation of TLR8 to promote FM inflammasome-dependent and inflammasome-independent mechanisms of inflammation. We propose that following signal one - TLR4 activation by LPS - the classic MyD88/NFκB pathway is initiated. While this may itself drive some of the FM inflammation, we propose that TLR4-mediated expression of miR-146a-3p subsequently triggers TLR8 activation which in turn contributes to FM IL-8 secretion and drives uric acid production - signal two. Uric acid-mediated NLRP3 inflammasome activation then results in FM IL-1β cleavage and secretion, amplifying FM inflammation and sustaining preterm birth. In summary, our studies have unveiled a role for TLR8-activating miR-146a-3p as a mediator of a FM chemotactic and inflammasome-mediated inflammatory responses to a bacterial insult, and thus may play a pathologic role in chorioamnionitis and subsequent preterm birth.

## Author Contribution

VMA and HMG designed the research. HG, CC and MT performed the experiments. HG and VMA analyzed the results and made the figures. HG and VMA drafted the paper. VMA, HMG, CC and MT revised the paper for important intellectual content.

## Acknowledgements

The authors would like to thank the staff of Labor and Delivery, Yale-New Haven Hospital and the Yale University Reproductive Sciences Biobank for blood and tissue collection. The authors would also like to thank Professor Richard Flavell for generously providing us with the *TLR7^−/−^/TLR8^−/−^*mice and for his guidance.

## Completing Interests

The authors have no competing interests.

