## Supplementary figures and images for "TLR8-Activating miR-146a-3p is an Intermediate Signal Contributing to Fetal Membrane Inflammation in Response to Bacterial LPS"

### Supplemental Figure 1

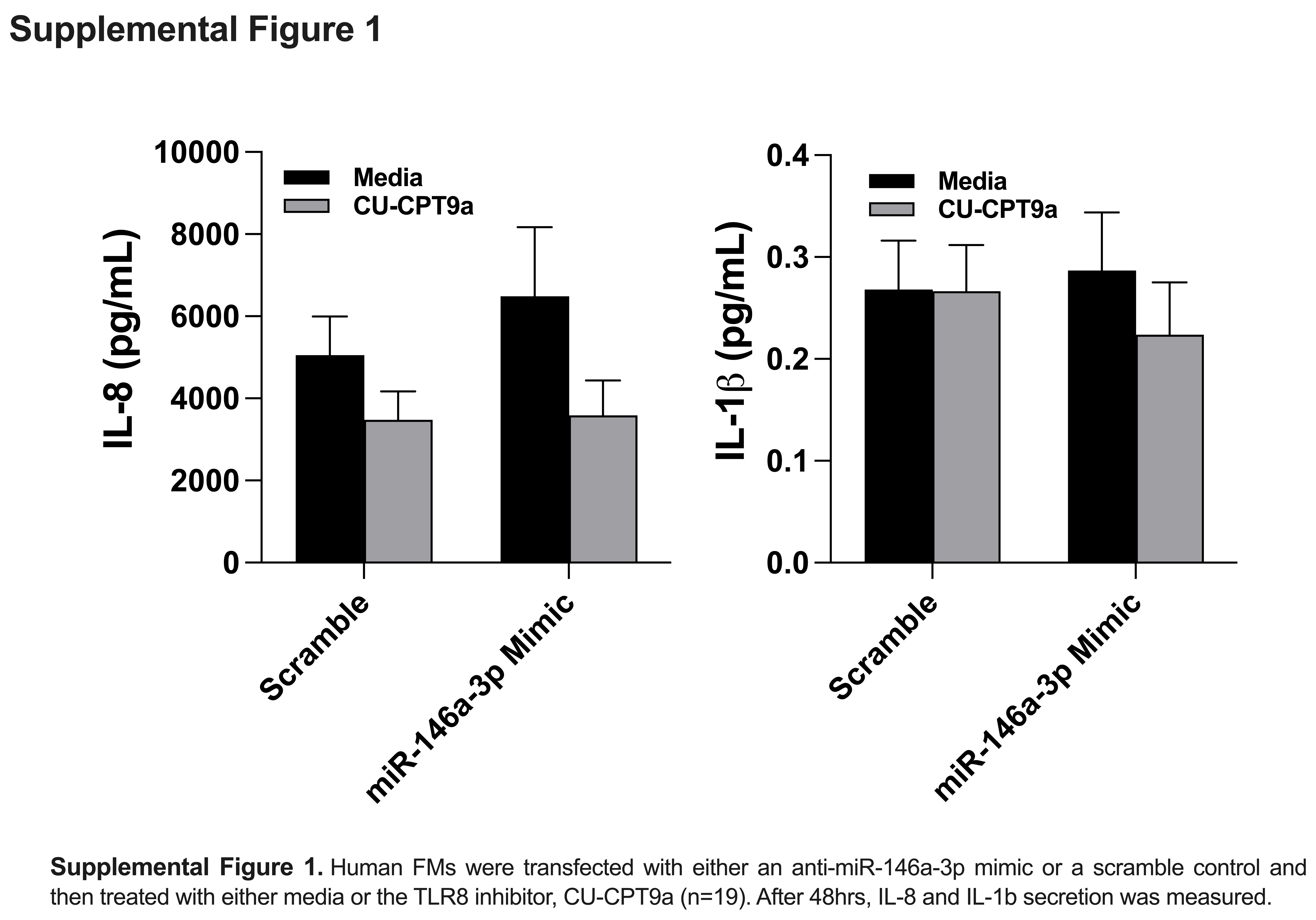
